# Instruction Alters the Influence of Allocentric Landmarks in a Reach Task

**DOI:** 10.1101/2024.04.11.589034

**Authors:** Lina Musa, Xiaogang Yan, J. Douglas Crawford

## Abstract

Allocentric landmarks have an implicit influence on aiming movements, but it is not clear how an explicit instruction (to aim relative to a landmark) influences reach accuracy and precision. Here, 12 participants performed a task with two instruction conditions (*egocentric* vs. *allocentric*), but with similar sensory and motor conditions. Participants fixated gaze near the centre of a display aligned with their right shoulder while a target stimulus briefly appeared alongside a visual landmark in one visual field. After a brief mask/memory delay the landmark then re-appeared at a different location (same or opposite visual field), creating an ego/allocentric conflict. In the *egocentric* condition, participants were instructed to ignore the landmark and point towards the remembered location of the target. In the *allocentric* condition, participants were instructed to remember the initial target location relative to the landmark and then reach relative to the shifted landmark (same/opposite visual field). To equalize motor execution between tasks, participants were instructed to anti-point (point to the visual field opposite to the remembered target) on 50 % of the egocentric trials. Participants were more accurate, precise, and quicker to react in the allocentric condition, especially when pointing to the opposite field. We also observed a visual field effect, where performance was worse overall in the right visual field. These results suggest that when egocentric and allocentric cues conflict, explicit use of the visual landmark provides better reach performance than reliance on noisy egocentric signals. Such instructions might aid rehabilitation when the egocentric system is compromised by disease or injury.

**Highlights:**

- 12 participants reached to remembered targets in the presence of a visual landmark
- Participants were instructed to ignore, or point relative to, the landmark
- The landmark instruction improved reaction time, precision, and accuracy
- These effects were stronger when pointing was cued toward the opposite visual field
- Knowledge of these rules might be used to enhance performance or in rehabilitation

## 1. Introduction

Humans use specific spatial reference frames to retain remembered visual information for a goal-directed action (Crawford et al., 2011; Soechting & Flanders, 1992). The visual system is thought to utilize two types of spatial reference frames, observer-centered, egocentric reference frames versus world-fixed, allocentric reference frames, often anchored relative to reliable landmarks (Byrne et al., 2010; Howard & Templeton, 1996; Vogeley & Fink, 2003). In ordinary circumstances, the brain integrates information from these two reference frames in Bayesian manner (Byrne & Crawford, 2010; Fiehler et al., 2014). However, certain circumstances may require one to ignore surrounding landmarks (i.e., when they are unstable or irrelevant to the task) or focus strongly on surrounding landmarks (such as a workspace fixed to a moving base). Various experiments have tapped into these mechanisms by instructing participants to use one reference frame over the other (Bryne & Crawford, 2010; Lemay et al., 2004). The question thus arises, which of these instructions leads to better performance?

Several studies have explored the implicit influence of visual landmarks on goal-directed actions such as saccades, reaches, or pointing towards a seen or remembered target. Previous findings have shown that reach targets can be remembered reasonably well in the absence of visual landmarks (Lemay & Stelmach, 2005; McIntyre et al., 1997; Vindras & Viviani, 1998) with certain stereotypical errors such as gaze-centered overshoots (Bock, 1986; Henriques et al., 1998). However, the addition of a visual landmark can influence reaching, reducing both constant and variable errors (Byrne et al., 2010; Krigolson & Heath, 2004; Lemay et al., 2004; Redon & Hay, 2005). The stabilizing influence of a landmark was particularly prominent in a task that involved remapping the reach target to the opposite visual hemifield, where one would expect egocentric signals to be less stable (Byrne et al., 2010). Conversely, the landmark had less stabilizing influence on behavior when they were shifted and rotated relative to the reach goal (Thaler & Todd, 2009). Finally, the addition of visual landmarks can negate the accumulation of reach errors after prolonged memory delays in the dark (Chen et al., 2011).

Normally, egocentric and allocentric cues agree, but they can also conflict, either in tasks that introduce egocentric noise or when the visual environment is unstable (Byrne & Crawford, 2010; Byrne et al., 2010, Chen et al., 2011). The latter has been replicated experimentally in cue-conflict tasks where the landmark is surreptitiously shifted relative to egocentric coordinates during a memory delay (Byrne & Crawford, 2010). In this situation, ego / allocentric cues appear to be optimally integrated, based on their relative reliability (Byrne & Crawford, 2010). Usually, more weight is placed on egocentric coordinates, such that the movement shifts approximately 1/3 in the direction of the landmark shift (Byrne & Crawford, 2010; Fiehler et al., 2014; Li et al., 2017). However, the specific weighting depends on task details. For example, Byrne & Crawford (2010) found that participants rely more on a landmark when it is perceived to be stable, or when gaze position was less stable. Further, in simulated naturalistic settings, landmarks had more influence when they were task-relevant, when more than one landmark was shifted in the same direction, and when the landmark was closer to the target (Fiehler et al., 2014). Thus, in the absence of explicit instructions, the visual system uses implicit algorithms to determine how to weight ego / allocentric cues. Recent physiological studies suggest that this implicit integration may occur in frontal cortex (Bharmauria et al., 2020, 2021).

Alternatively, people can be instructed to ignore or put full weight on a visual landmark, as in common driving instructions like ‘ignore the first stop sign and turn right at the second’. Likewise, experimental participants can be instructed to either ignore a landmark or reach to a fixed egocentric location relative to the landmark. Such instructions were used in neuropsychological and neuroimaging experiment that suggesting involvement of the ventral visual stream in allocentric representations and dorsal stream in egocentric transformations (Chen et al., 2014, 2018, Goodale et al., 2004; Schenk, 2006). However, the influence of these instructions on behavior is less clear. For example, in their control tasks, Byrne & Crawford (2010) found no overall difference in performance when participants were explicitly instructed to use egocentric or allocentric cues, but there was no cue-conflict in these tasks and the visual stimuli were different. Thus, it remains unclear what effect the instruction itself (to prioritize egocentric vs. allocentric cues) has on behavior, particularly when these two cues conflict.

Here, we tested the influence of instruction (to use or not use a landmark) in a cue-conflict, memory-guided reach task where the visual stimuli were equal and balanced across tasks. In this way we could isolate and directly compare how top-down reliance on allocentric vs. egocentric cues affected reach behavior. It has been suggested that landmark-centered encoding is a less noisy process and more stable over a time delay (Byrne et al., 2010; Chen et al., 2011). Thus, we predicted that instruction to attend to and use the landmark for spatial coding would increase weighting on the more stable code, and therefore improve performance, especially when the egocentric noise is higher (Byrne & Crawford, 2010, Chen et al., 2011). To simulate the latter case, we included an ‘anti-reach’ condition where participants had to reach to the opposite visual field as the viewed target. We found that 1) reaching was less variable and more accurate in the allocentric instruction tasks than egocentric instruction tasks (especially in the right visual field) and 2) the beneficial effect of allocentric encoding was more pronounced when participants were required to respond in the visual field opposite to stimulus encoding.

## 2. Material and Methods

### 2.1 Participants

13 individuals (7 males and 6 females; ages 20 – 33) provided informed consent to participate in this study. All participantswereright-handed with no neuromuscular or uncorrected visualdeficits, based on self-report. Data from one participant was excluded from further analysis because they did not meet the inclusion criteria described below, leaving 12 participants for data analysis. This met the sample size required for sufficient power (see §1.5. *Sample Size Analysis*). All participants were naïve to the purpose of the experiment and given monetary compensation for their time. The experimental procedures were approved by the York Human Participants Review Subcommittee.

### 2.2 Apparatus

The experiment was conducted in darkness and participants wore a black glove on their right hand to avoid any ambient reflections off their right hand. Participants sat behind a table, on a chair of an adjustable height (Fig. 1). The head was stabilized on a personalized bite bar made of a dental impression compound (Kerr Corporation, Orange, CA). Participants’ right hand was positioned on a button box on the table-top, directly in front of them. The button box was used to control the pace of the experiment and was the designated start position for reaching. A customized ring with a 3 x 3 array of Infrared-emitting diodes (IREDs) was attached to the participants’ right index finger and their 3D-position was continuously recorded by two OptoTrack 3020 tracking systems (Northern Digital, Waterloo, Ontario, Canada). Gaze direction was monitored from only the right eye, through the EyeLink II infrared eye-tracking system, (SR Research, Mississauga, Ontario, Canada), that was mounted to the bite bar stand. The stimulus display (described below) was presented 50 cm ahead of the eye and 152 mm to the right / 91 mm bellow the centre of the bite bar stand, approximately aligned with the right shoulder bone (acromion). The height of the chair was adjusted so that the acromion was aligned with the centre of the display panel. This arrangement ensured that stimuli were centred in the right arm’s mechanical range and within easy reach. During reaches participants’ head was facing forward, but gaze was fixated at the centre of this display, such that shoulder and central visual coordinates aligned. Audio instructions were delivered from a speaker. Two 40-watt desk lamps, placed on either side of the desk, were turned on every 10 trials (2 min), to eliminate potential dark adaptation. The experimental set-up is shown in Fig. 1.

**Fig. 1:**
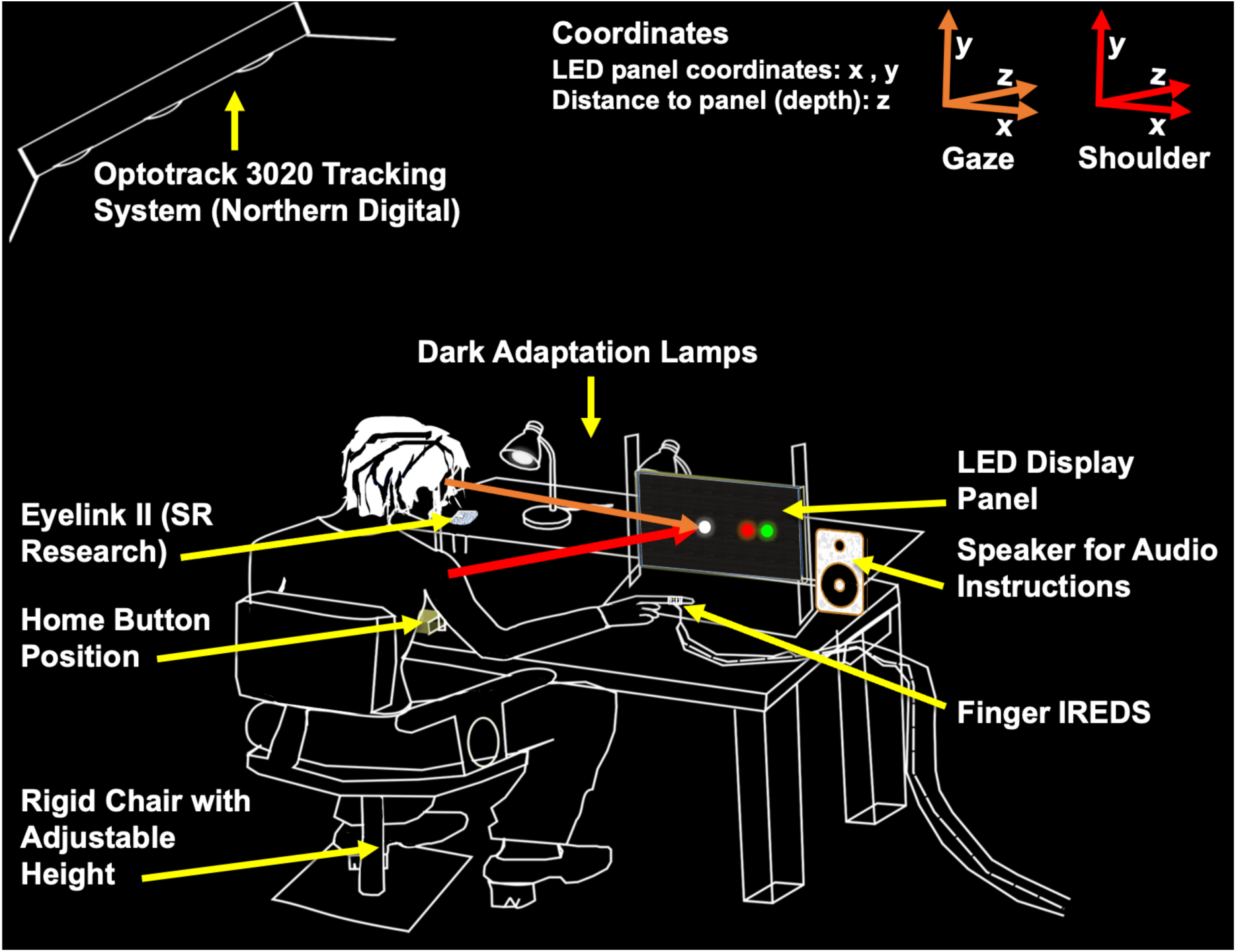
Experimental set-up. From left to right and top to bottom: OptoTrack 3020 tracking systems (Northern Digital, Waterloo, Ontario, Canada), on either side of the room (second one not shown in the figure) were used to track the finger motion. Eyelink II cameras (SR Research, Mississauga, Ontario, Canada) were dismounted from the headset and installed on the bite-bar stand. The left camera is occluded in this view of the set-up. Home button box (yellow in the figure), on the desk immediately in front of participants was used to control the pace of the experiment. Participants’ height was adjusted relative to the fixed bite-bar height using a metallic screw on a rigid chair with an adjustable height. Two 40 W dark adaptation desk lamps illuminated the dark room during breaks and every 3 trials. A custom made 307 mm x 161 mm wooden panel, fitted with light emitting diodes (LEDs) was used to display stimuli. Audio instructions were played from two desktop speakers, one of the speakers is not shown in the figure. The finger pointing device was a customized ring with a 3x3 array of Infrared-emitting diodes (IREDs) that continuously relayed signal to the OptoTrack 3020 tracking systems.

### 2.3 Calibration

Before each experiment, participants completed a set of calibration procedures. Eye position calibration was done through sequential fixation of 5 light emitting diodes (LEDs) including the centre position and the four corners of the display panel (See Fig. 2A). They then completed two OptoTrak calibration sessions. Finger-tip position was calibrated by having participants point with a pre-calibrated ‘cross’ of IREDs on a rigid body fixed to the right index finger. Further calibration was done by having participants successively point to the 4 corner positions in the LED display. IRED position data in the OptoTrak intrinsic coordinate system was compared offline with known positions of the calibration dot positions to create a linear mapping between IRED positions and the screen coordinates. This procedure allowed for the conversion of recorded finger-tip position into screen and then visual coordinates, as described below.

**Fig. 2:**
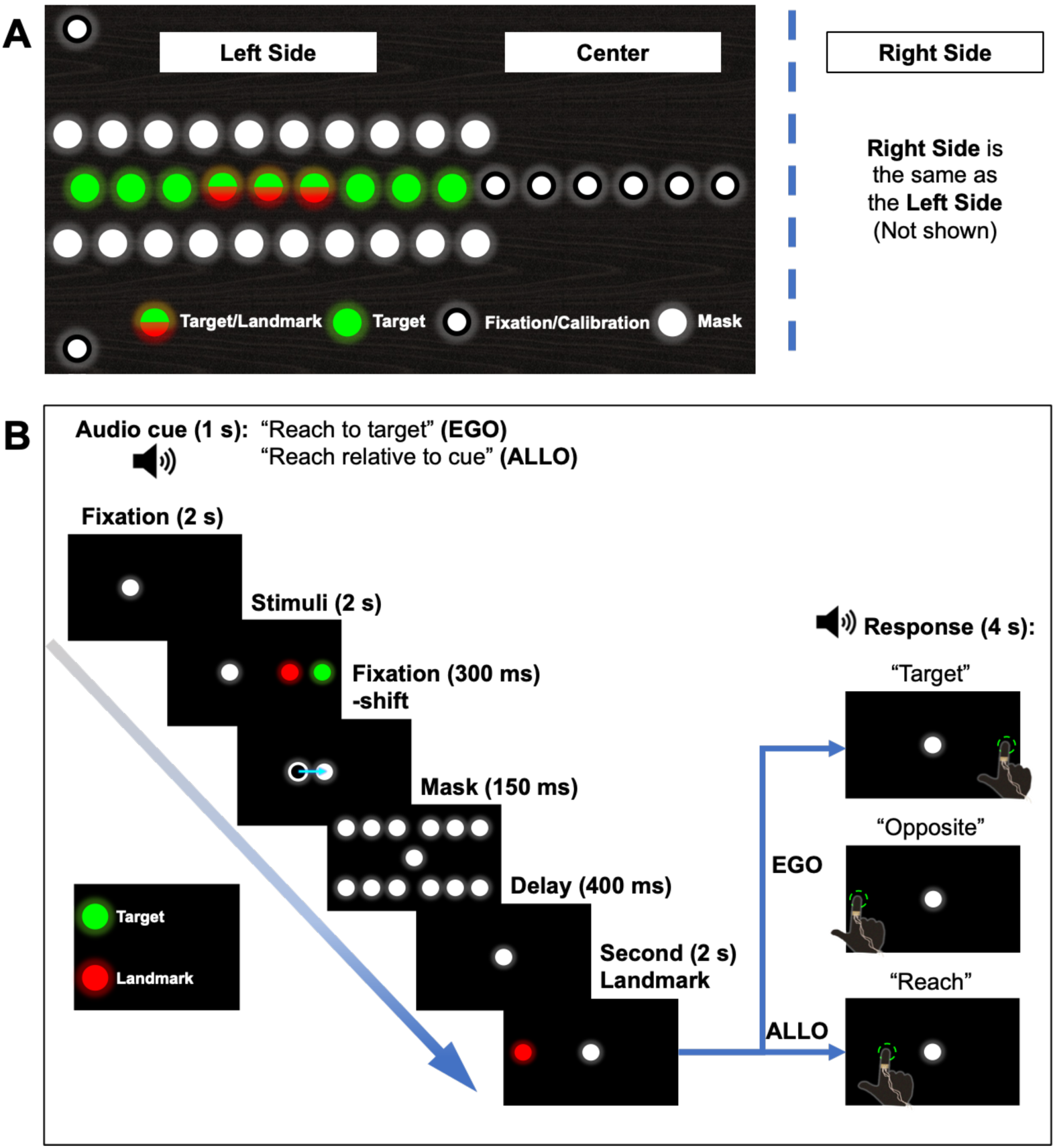
Experimental Stimuli and Paradigm. **(A)** Stimuli. The stimuli were displayed in a horizontal array. Left side and center of the array are shown, while the right side is the same as the left side (mirror image). The central fixation, off-centre fixation locations and eye calibration points were displayed by white LEDs. The green LED target was randomly displayed in one of the 18 (9 left, 9 right) LED positions. The position of the first target LED was 3 circles (4.57 degrees) from the screen centre. The red landmark simultaneously appeared on the same side of the screen as the target, displayed by one of the 3 LEDs in the middle of the 9 target LEDs (half red, half green circles in the figure), while the second red landmark randomly appeared in one of the remaining landmark positions. **(B)** Paradigm. The order of a typical trial is shown in the figure. Each trial began with an audio instruction, where participants were instructed to remember the spatial location of the target, or the position of the target relative to the landmark. The response audio depended on task; *EGO* trials were followed by “Target”, instructing participants to point toward the target, or “Opposite”, instructing participants to point to the mirror opposite side, while all *ALLO* trials were simply followed by “Reach”, instructing participants to point to the remembered target position relative to the second landmark.

### 2.4 Visual Stimuli and Basic Task

The experimental paradigm was based on a simpler paradigm used previously in an fMRI experiment (Chen et al., 2014). The task involved touching the remembered location of a transient visual target (that always appeared simultaneously with a visual landmark) as accurately as possible. Fig. 2A shows the visual/touch display. LEDs (1.15 degrees light emitting diodes; LEDs) were fixed to a wooden panel (307 mm x 161 mm). Target, landmark and fixation LEDs were placed at 1 cm (∼1.15 visual degrees visual angle) intervals along this panel. Seven white gaze fixation LEDs were placed at centre and 1.15, 2.29, 3.43 degrees to the left and right of the centre. 18 green reach target LEDs were placed in the periphery, starting at a 4.57 degrees distance from the centre. 6 red landmark LEDs were positioned 7.96, 9.09 or 11.31 degrees, to the right or left of the centre. 40 LEDs used as a visual mask were placed 1.15 degrees above and below the target/landmark stimuli, in-between the target/landmark LED positions.

The same stimulus paradigm was used in all experiments, with LED positions and task instructions randomized as described below. The sequence of stimulus events is shown in Fig. 2B. Participants started seated comfortably with their hand placed on the button box near their chest. At the beginning of each trial, they received an audio instruction from the speakers telling them how to reach (see next section for details). Participants were then required to visually fixate one of the 6 non-centre white LEDs. After 2 s a green target LED was displayed to the right or the left of the centre. This was accompanied simultaneously by presentation of one of the 6 red landmark LEDs relative to the target, on the same side of the centre as the target. After 2 s, both the target and landmark stimuli disappeared and the gaze fixation point moved to centre, allowing 300 ms for a saccade to that position. This was done so that the fixation LED could not be used itself as a reliable allocentric landmark (Chen et al., 2014). The array of white ‘mask’ LEDs was then illuminated for 150 ms to negate visual afterimages and focus memory resources (Medendorp et al., 2003). After the mask, the centre fixation LED reappeared, requiring continued central fixation for the rest of the trial. A landmark then reappeared for 2 s, but always at a different location than it was initially viewed, creating a conflict between the location of the target in egocentric coordinates versus allocentric coordinates. At this point participants were instructed to reach via the speakers, and then were given 4 s to touch the stimulus panel with the right index finger as accurately as possible, based on the instruction they had received at the start of the trial. They then returned their hand to the starting position and pressed the button when they were ready for another trial.

### 2.5 Task Instruction and Conditions

As noted above, the visual paradigms used in this task were always the same for different task conditions, what differed between conditions was the task instruction delivered via the speakers at the start of the trial. In the *Allocentric (ALLO) Condition,* participants were instructed to “*Reach relative to cue*”, i.e., they were required to remember the position of the green target, relative to the red landmark. Since the landmark was shifted to a new position at this time, egocentric coordinates provided no useful cues for this task. In this case, the instruction to reach at the end was simply ‘*reach*’. In contrast, in the *Egocentric (EGO) Condition*, participants heard “*Reach to target”* at the beginning of the task. In this condition, participants were instructed to ignore the landmark (which in this condition provides invalid information). Henceforth, we refer to these as the *EGO Instruction* and the *ALLO Instruction* conditions.

To further challenge the system, in half of the trials participants were required to perform ‘anti-reaches’ to the visual field opposite to the field where they saw the target (Cappadocia et al., 2017; Gail & Andersen, 2006). In the *ALLO Condition*, this was done by shifting the landmark to the opposite hemifield. In the *EGO condition*, an additional instruction was provided. When participates heard ‘*target*’, they reached toward the original location of the target (‘*PRO-Task*’). When participants heard ‘*opposite*’ they were expected to reach toward the mirror opposite position, relative to the screen midline (‘*ANTI Task’*). To be consistent with literature, we then divided our data *into EGO PRO/ANTI -* or *ALLO PRO/ANTI -reach* trials (based on the final reach direction relative to initial target). Finally, in both *PRO* and *ANTI* trials, stimuli were arranged so *EGO/ALLO* goals covered the same range.

### 2.6 Experimental Design

Before each experiment, participants engaged in a practice session to ensure that they understood the instructions and were able to perform each element of the task correctly. This consisted of 3 practice blocks of 10 trials. The first two practice blocks were made up of each *EGO/ALLO Instruction* condition separately and the last practice block was randomly interleaved. After they were successful and clear on the instructions, they proceeded to the actual experiment. Actual experiments were divided into 3 blocks of 72 trials. Participants had a 5-min rest period between blocks and four 2-min breaks within each block (with room lights fully illuminated). Individual trials lasted 12.05 s in both instruction conditions, but the experiment was self-paced, subsequent trials were initiated through a button press. On average, participants took 21.65 mins to complete an entire block. The order of instructions, target locations, and landmark locations was pseudorandomized beforehand so that they were unpredictable for the participants but provided an equal dataset with distributed stimuli for each experimental condition.

### 2.7 Data Analysis

All data obtained from OptoTrak and EyeLink were analysed offline using a custom software written in MatLab R2019a (Math Works). A program was written to generate a mapping between the Optotrak coordinates and the position of the fingers in screen coordinates of the fingertip. This was done by utilizing the data exported from the screen calibration session and known screen coordinates of the 4 calibration points on the screen corners. This mapping was then utilized to obtain reach endpoints in screen coordinates for analysis. Finally, target and touch positions in screen coordinates were converted into visual angle (relative to the central fixation point) using geometric measures obtained from the laboratory set-up.

#### 2.7.1 Exclusion Criteria

Movement kinematics were inspected to ascertain that the participants followed instructions and a program was written to automatically average eye fixation, from the period of the second landmark to the go signal. Trials were excluded if eye variation was greater than 2 degrees in the horizontal direction for more than 20 % of time. Trials with brief saccades to remembered target location that then returned were not excluded, because this does not appear to influence pointing accuracy (Van Pelt & Medendorp, 2007). However, trials where participants looked at the landmark were excluded. Finally, trials that included anticipatory reaches (reaches initiated before the audio instructions) were excluded from further analysis.

Overall, 12 participants who had at least 175 viable trials (81.02 %) were included in the study. Participants completed on average (± standard deviation) 72.90 ± 20.58 *EGO Instruction* trials, on average 37.80 ± 8.98 of trials were trials in which the target was viewed, and the movement was executed in the same visual field while 36.70 ± 10.60 of trials were trials in which the movement was executed in the opposite visual field. On average, participants also completed 70.60 ± 18.53 *ALLO Instruction* trials, 32.70 ± 7.81 of trials were on the same visual field and 35.70 ± 12.99 of trials were in the opposite visual field.

#### 2.7.2 Kinematic Analysis

Using a customized MatLab program, movement start was marked at 20 % maximum resultant velocity (Vmax), while movement end was marked at 8 % of Vmax. Reach endpoint was found by averaging 30 % of points at 8 % Vmax and at 1.15 times the minimum finger to screen distance (distance in the z direction). The screen relative coordinates of reach endpoints in the horizontal (x) and vertical (y) dimensions were used to compute the variable error, a measure of the distance of reach endpoints from the mean final position. R 4.1.2 (Core Team, 2021) was used to create 95 % confidence ellipse of the scatter of reach endpoints. The area of the ellipse was found by first computing eigenvalues (σ1, σ2) of the covariance matrix of reach endpoints. The eigenvalues were then used to derive half the lengths of the semi-major (principal axis) and semi-minor (orthogonal to the principal axis) axes of the 95 % confidence ellipse and used to compute the ellipse area (formula [1] shown below). The ellipse areas were then used to compare the *EGO* and *ALLO Instruction* data. Mean ellipses for each target were constructed by averaging the covariance matrices of the corresponding ellipses of individual participants.

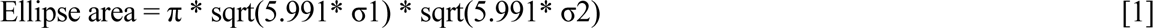

The overshoot error in horizontal reach endpoints was used to analyse response accuracy. The horizontal overshoot error in reaching response at each target location was found by subtracting the difference between the reach endpoint to screen midpoint distance, and the expected target location to the screen midpoint distance. The expected target location was the same as the initial location for *EGO/PRO* reaches, mirror opposite to the target location for *EGO/ANTI* reaches but based on the shifted landmark for all allocentric reaches. The mean overshoot error was found by averaging the magnitude and direction of the reaching error (positive to the right, negative to the left), separately for each instruction condition (*EGO/ALLO*), task condition (*PRO/ANTI*), and visual field of response. Ellipse areas, reaction times and movement times were averaged in a similar manner. The reaction time was the period between the go signal and the movement start, while the movement time was the period between the movement start and the movement end.

### 2.8 Sample Size Analysis

Sample size was calculated based on the simulation approach described by Green & MacLeod, (2016), using the simr R package (Core Team, 2021), which calculates the power for generalized linear mixed models from the lme4 R package (Core Team, 2021). The power calculations were based on Monte Carlo simulations. Reaching variance was obtained from a pilot study of 2 individuals and 72 observations for each (*EGO/ALLO*) instruction condition. A mixed-effects model was fitted on pilot data, to obtain an estimated effect size of 0.267 for reaching variance. Using the obtained effect size and the simulation package, the sample size was gradually increased to achieve a power of 85 %. The sample size required to achieve this power was 12.

### 2.9 Statistical Analysis

Linear mixed effects models were used to analyse differences in ellipse areas, overshoot errors, reaction times and movement times, using the lme4 R package (Core Team, 2021). The reaching responses relative to the expected target positions were fit with quadratic mixed effects models, using lme4 R package (Core Team, 2021), to determine differences in reaching behaviour due to the task conditions. Interaction contrasts were performed using the eemeans R package (Core Team, 2021).

For statistical analysis, we grouped our data into 4 task conditions, as follows:

1. *EGO Instruction / PRO Task*
2. *EGO Instruction / ANTI Task*
3. *ALLO Instruction / PRO Task*
4. *ALLO Instruction / ANTI Task*

In addition, based on the findings of Byrne et al. (2010), we also sorted these data into hemifields (*LEFT / RIGHT*), based on the final direction of the instructed reach goals.

## 3. Results

Our experiment was designed to test if an explicit instruction to reach relative to an allocentric visual cue (as opposed to ignoring this cue and relying on egocentric coordinates) would alter reach performance in the presence of otherwise identical stimuli. For example, the top panel (A) of Figure 3 represents the case where both the target and landmark were presented in the right visual field, and after the target disappeared the landmark shifted to the left visual field. The bottom two panels of Figure 3 (B) each display 6 representative hand trajectories (black lines) and gaze trajectories (grey lines) associated with the stimuli in the top panel. Both panels show *ANTI-Task* trials but in (B1) the participant was given the egocentric ‘opposite’ instruction whereas in the other case (B2) the participant was instructed to point relative to the (shifted) landmark. As one can see, the participant was able to perform both tasks, although with subtle differences. In total, 12 (of 13) participants completed a sufficient number of these trials in all versions of our task (including the easier *PRO-task* versions) for statistical analysis of their data. An analysis focusing on comparisons of precision, accuracy, and reaction time between conditions will follow.

**Fig. 3:**
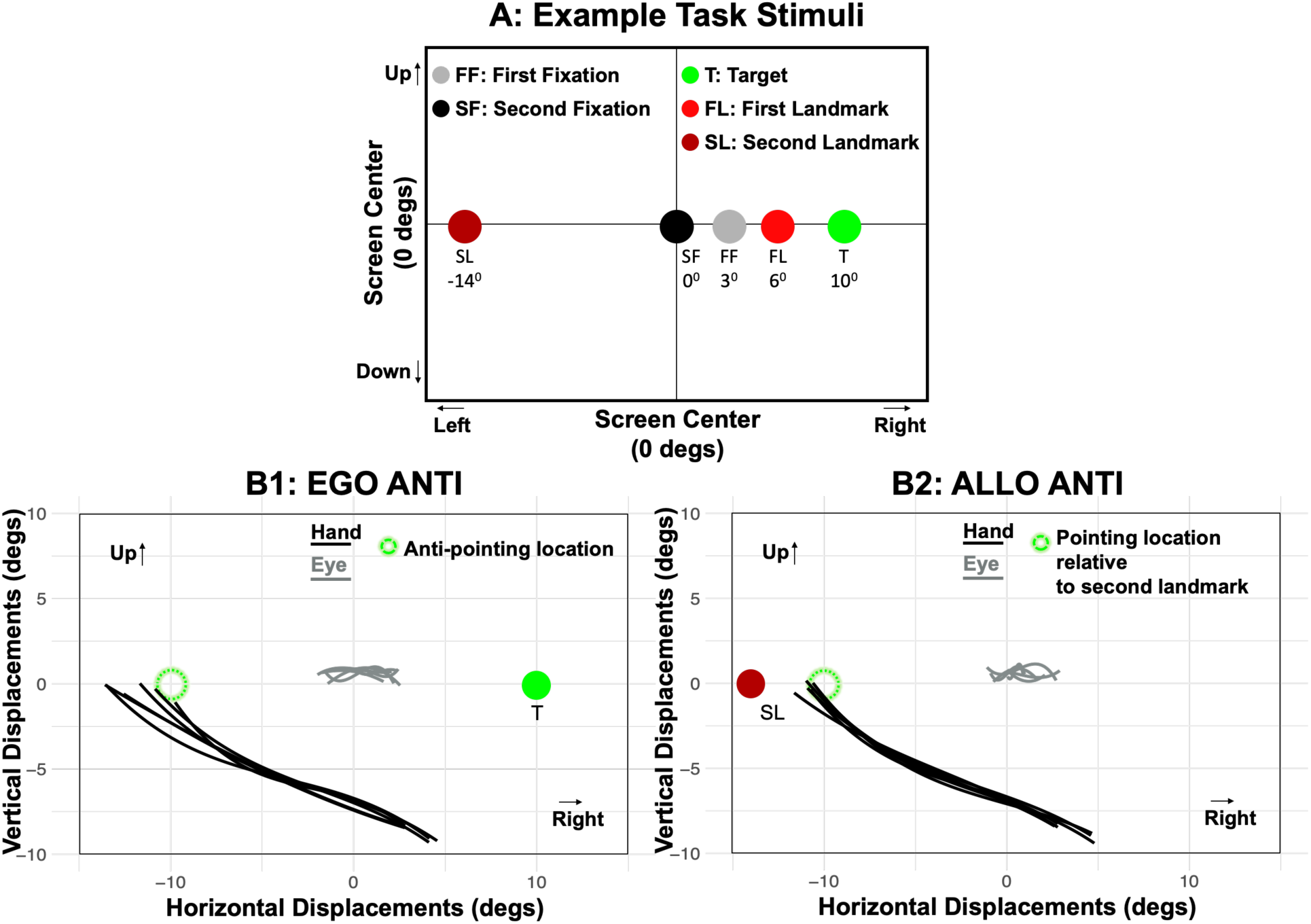
Typical Eye and Finger Trajectories. **(A)** Spatial location of example stimuli (see graphic key for details). **(B1)** *EGO / ANTI condition*, where the participant was instructed to touch mirror opposite spatial location of the remembered target position. The same color conventions as A are used, except the unfilled green circle shows the ideal goal position, the black lines show the 2D finger trajectory for 6 example trials, and the grey lines show the corresponding 2D gaze locations during central fixation, where gaze was required to remain within +/- 2 degrees of the fixation point 80 % of the time. **(B2)** *ALLO / ANTI condition*, where the landmark appeared in the opposite side of the fixation (same graphic conventions as B1).

### 3.1 Reach Accuracy and precision: General Observations

Fig. 4 provides a spatial overview of our pointing results in all four conditions, plotted in visual coordinates. The top portion of each panel shows the labeled 2-D target positions (black dots), corresponding to reach data points (medium circles), and their 95 % confidence ellipses (large ovals), using the same color code for each (light shade near centre, darker toward periphery). The lower parts of each panel show the corresponding 1-D probability densities for each target position along the horizontal dimension (the dimension in which targets varied). The leftward panels (A & B) exhibit the plots for a single participant, while the rightward panels (C & D) portray the mean ellipses and probability densities across participants. The plots pertaining to the *EGO Instruction* (blue) are showcased in blue on the top panels (A & C), whereas those concerning the *ALLO Instruction* (red) are presented in red on the bottom panels (B & D).

**Fig. 4:**
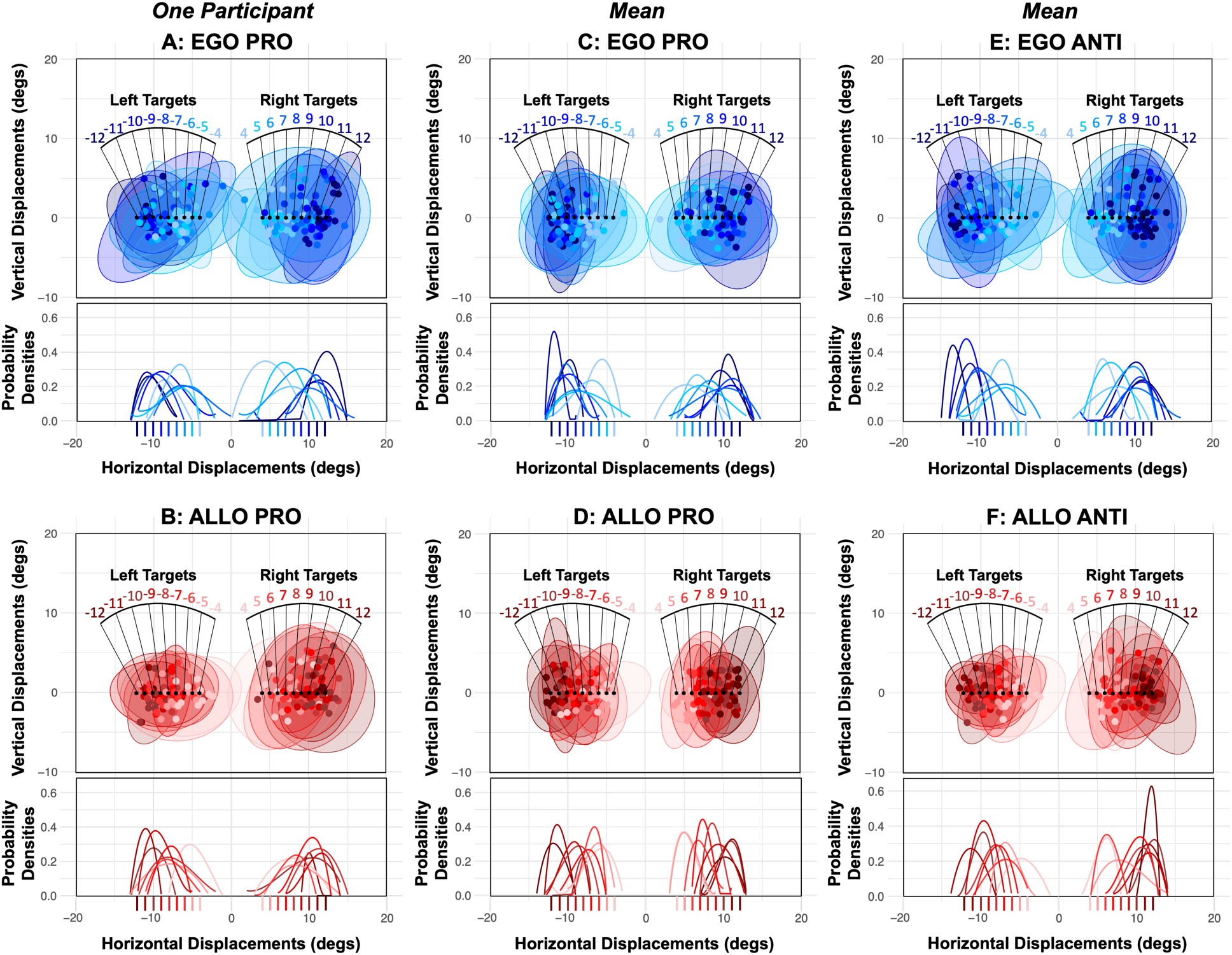
Reach Endpoint Variability. Panels for a representative participant **(A)** & **(B):** 95 % confidence ellipses (2D reach endpoint variability) are shown at the top of each panel and horizontal reach endpoints probability densities (1D reach endpoint variability) are shown at the bottom of each panel. The small, filled circles are individual reach endpoints relative to the central fixation. Ellipses, probability density curves and reach endpoints are all color coded by initial target spatial location. The color gradient, from light to dark, represents targets that are central to peripheral. Panels **(C)**, **(D), (E) & (F):** Mean ellipses, mean horizontal probability densities and mean reach endpoints of 12 participants. Panels **(A) & (C):** *PRO-Task, EGO Instruction* (top) in blue and panels **(B) & (D):** *PRO-Task, ALLO Instruction* (bottom) in red. **(E):** *ANTI-Task, EGO Instruction* (top) in blue and panels. **(F):** *ANTI-Task, ALLO Instruction* (top) in red and panels. The target positions are referenced using black dots in the ellipse figures (the exact distances are labelled on the plot) and as axis ticks on the x-axis in the 1-D density plot, both target position labels are color-coded using the same color gradient.

Some qualitative observations are that 1) the distribution of error was relatively large, as one might expect in a memory-guided task, 2) the vertical distributions tended to be larger than the horizontal distribution, and 3), the distributions did not overlap, but rather shifted with target location, and 4) the *ALLO Instruction* distributions seem slightly larger in the *PRO-Task* conditions. Fig 4 right column, panels (E & F), illustrate the average ellipses for the *ANTI -task* data using the same conventions as the middle column of Fig. 4. Here, the *EGO* distributions were still larger, and thus the difference between the *EGO* and *ALLO* distributions seems clearer, at least in the left visual field, where the *ALLO* distributions are visibly smaller than the *EGO* distributions. Each of these observations will be quantified in more detail below.

### 3.2 Quantification of Reaching Variance

Fig. 5 quantifies the distributions of the ellipse fit areas (averaged within each participant) for the left and right visual field targets, with *EGO Instruction* data depicted in blue and *ALLO* in red. The *PRO Task* data are depicted in the top panel (A), while the *ANTI task* data are depicted in the bottom panel (B). As one can see, the general trend is for smaller ellipses fits (i.e., higher precision) in *ALLO instruction* data. To quantify these data, we performed a statistical analysis of the instruction influence (*EGO/ALLO*), task (*PRO/ANTI*) and visual field (*LEFT/RIGHT*) ellipse fits using a generalized linear mixed model. The analysis included the interaction of spatial instructions with both task and visual field of response. No significant three-way interaction effect was found, leading to the fitting of a simpler model (Table 1).

**Fig. 5:**
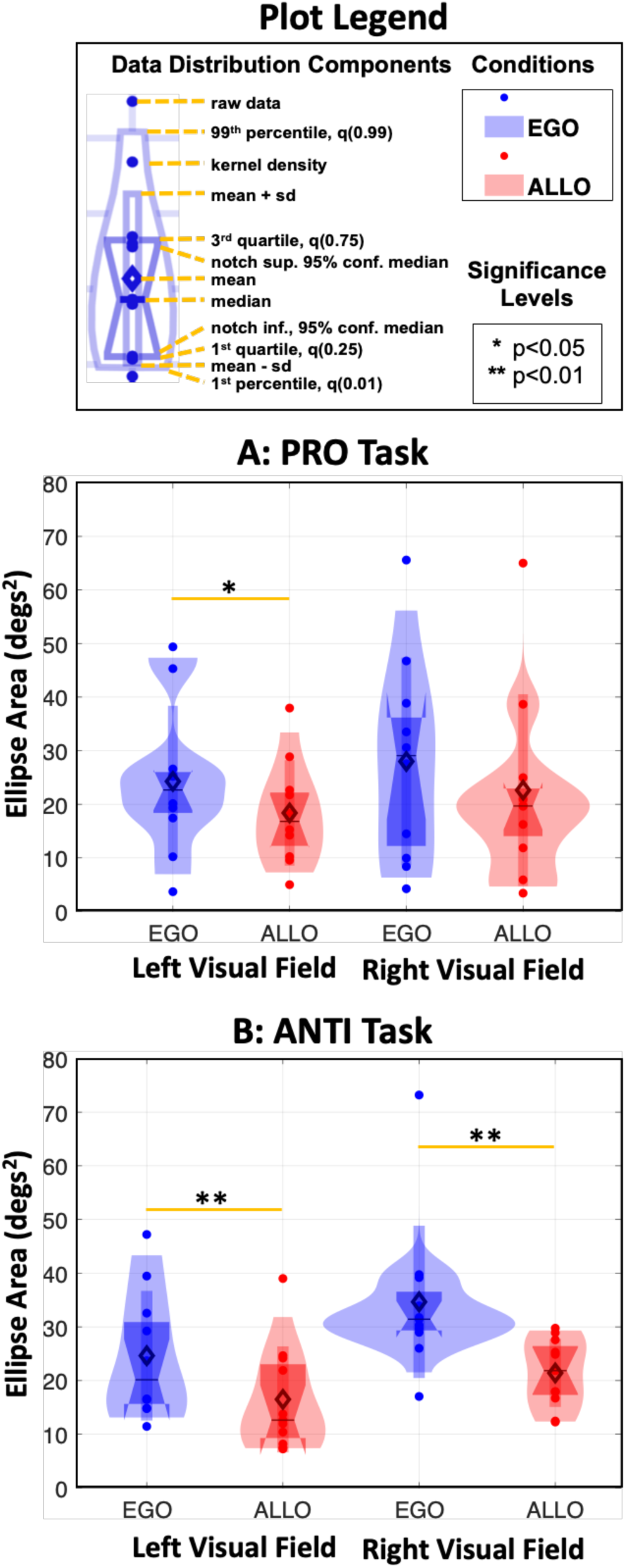
Reach Endpoint Ellipse Areas. Violin Plots of ellipse areas averaged over right/left visual field targets, and for each participant. The legend on top explains the conventions used to generate these plots. Each plot (A/B) summarises the observed trends in the left visual field (left side) and right visual field (right) for each task/instruction condition. **(A)** Mean areas of 95 % confidence ellipses for the *PRO-Task* data. **(B)** Mean areas of 95 % confidence ellipses for the *ANTI Task* data. *EGO* and *ALLO Instruction* conditions are shown in the figure as blue and red, respectively. Significant differences are shown by an asterisk, *p < 0.05, **p < 0.01.

**Table 1.** Fixed Effects of Multi-level Model using Ellipse Areas as the Criterion.

| Predictor | *B* | *B*<br>95 % CI<br>[LL, UL] | *t value* | *p* |
| --- | --- | --- | --- | --- |
| (Intercept) | 16.67** | [9.70, 23.61] | 4.77 | 8.5 e -6** |
| Task (*PRO*) | 1.55 | [-6.62, 9.71] | 0.377 | 0.71 |
| Instruction (*EGO*) | -5.65 ** | [-6.37, -4.92] | -11.57 | 2.0 e -15** |
| Visual Field (RIGHT) | 4.60 | [-3.56, 12.75] | 1.12 | 0.26 |
| Task<br>(*PRO*) x Instruction<br>(*EGO*) | -1.65** | [-2.37, -0.92] | -4.44 | 1.0 e -5** |
| Visual Field (RIGHT) x<br>Instruction (*EGO*) | 2.21 | [-9.33, 13.74] | 0.38 | 0.71 |

*Note.* *B* represents standardized regression weights. *LL* and *UL* indicate the lower and upper limits of the confidence interval, respectively. Model Adjusted $R^2 = 0.145^*$ , 95 % CI [0.00,0.24]
\* indicates $p < 0.05$ . \*\* indicates $p < .01$ .

The outcome of this analysis revealed a significant effect of spatial instruction, with ellipse areas in the *EGO Instruction* condition (mean ± SD, 29.87 ± 4.66 degrees^2^) being significantly larger i.e., less precise, than the *ALLO Instruction* condition (20.32 ± 3.77 degrees^2^) (p = 2.0 e -15). The *ALLO Instruction* led to improved precision consistently across visual fields of response, especially when participants had to point opposite to the visual field of response. Post-hoc interaction contrasts, averaged across visual fields, showed that participants were significantly less precise in the *EGO* than the *ALLO Instruction* condition, especially when they had to point opposite to the visual field of response (p = 1.0 e -5).

### 3.2 Reaching Accuracy

As noted above, the horizontal positions of the response distributions tended to vary with corresponding desired target locations (Fig. 4), suggesting that participants did not simply point to the left or right visual field. To quantify the accuracy of these responses, the absolute horizontal values of participants’ end points (averaged across trials for each target within each participant) were regressed relative to the corresponding ideal target locations (Fig. 6). A general observation is that the distribution of the *ALLO Instruction* data (red) appears to be more compact and follows the line of unity (ideal accuracy) more closely that the *EGO Instruction* data (blue).

**Fig. 6.**
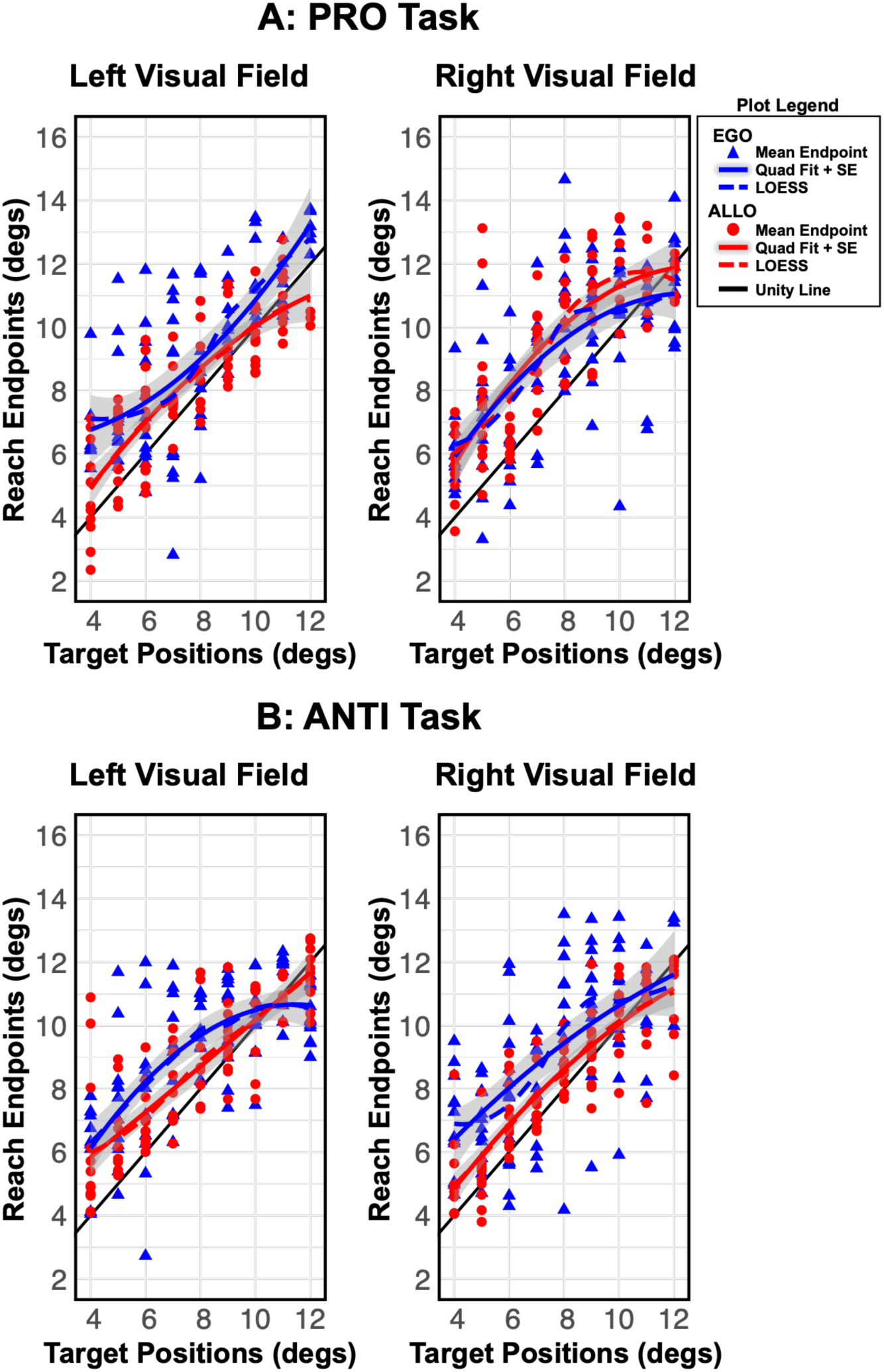
Scatterplots of the Horizontal component of Reach Endpoints vs Expected Target Positions. See graphic key (upper right) for details. The scatter plots of participants’ horizontal reach endpoints in the right visual field (right panel) and left visual field (left panel) were fitted with a quadratic line (solid coloured line). **(A)** In the *PRO Task* data. **(B)** In the *ANTI-Task* data. The *EGO* and *ALLO Instruction* conditions are shown in blue and red, respectively. The dashed colored lines are the locally estimated scatterplot smoothing (LOESS) lines. Black dashed line in figures is the line of unity. Shaded grey area shows the standard error of the estimate.

To select the most appropriate model to quantify these data, we employed a k-fold (leave-one-out) cross-validation, comparing the fit of three models: a linear regression model, a quadratic regression model centered at the first target, and a model with nominal predictors. The quadratic model exhibited a significantly better fit, as indicated by a lower root-mean-squared-error (RMSE) and a higher R-squared value. These fits are shown as curved lines, color coded lines in Fig. 6.

Table 2 summarizes the relative contributions (β) of the model parameters, indicating those which were significant (*), and their 95 % confidence intervals (CI). Overall, this fit yielded a significant (p < 0.01) R^2^ of 0.534 [0.48, 0.57]. The relationship between the participants’ reach endpoints and the targets demonstrated a significant negative quadratic trend (P = 0.02), suggesting a decrease in slope for more peripheral targets, possibly due to the decreased perceptual acuity of peripheral targets (see ‘Overshoot Errors’ below). This effect was significantly more pronounced in the *EGO Instruction* Condition (p = 4.0 e-3). In the *EGO instruction* condition, participants exhibited a significant decrease in the slope between reach endpoints and target location, indicating more accuracy in the *ALLO Instruction* condition. Reaching trajectory was not significantly influenced by the task (*PRO/ANTI)*, but a significant visual field effect was observed (P = 8.9 e-3), where participants exhibited a more negative quadratic trend and an increase in the tangential linear slope at the 4 degrees target. This suggests that participants’ reaches in the right visual field were relatively accurate for more central targets but demonstrated poor reaching performance for peripheral targets.

**Table 2.**
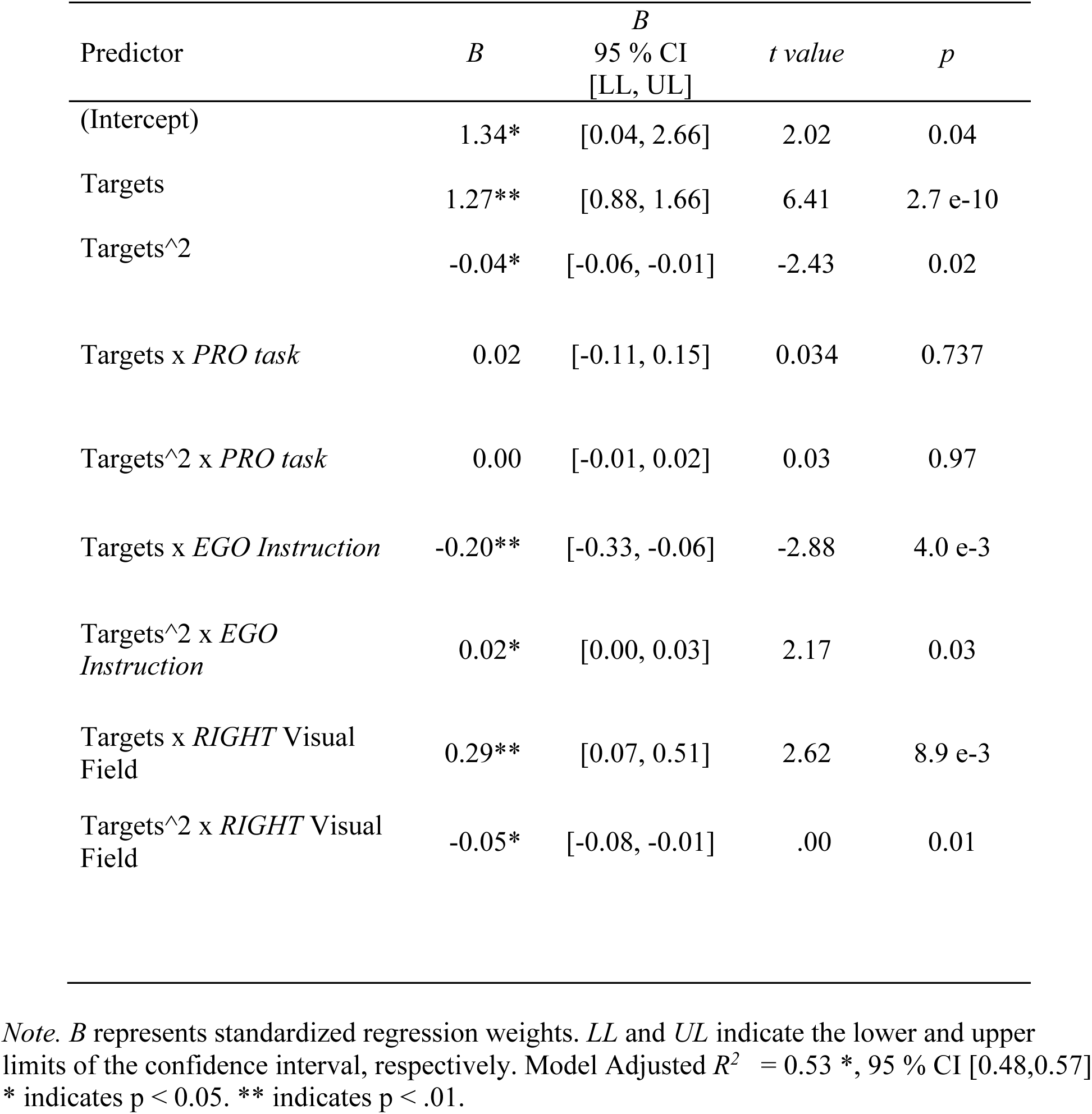
Fixed Effects of Multi-level Quadratic Model using Reach Endpoints as the Criterion.

### 3.4 Influence of Landmark Proximity and Shift Distance on Reach Endpoints

As described above, reach precision and accuracy were influenced by the EGO / ALLO instruction, but there was an influence of specific stimulus parameters. Specifically, the initial distance between the landmark and target (1-3 degrees), and the amplitude of the landmark shift (1-3 degrees for the *PRO-Task*, 9-14 degrees for the *ANTI-task*). If participants were able to suppress landmark influence after the *EGO-Instruction*, one would expect these parameters to have no effect, whereas they might have an influence after the *ALLO-Instruction*. To test this, two mixed-effects models were employed separately on *EGO-Instruction* and *ALLO-Instruction* conditions to assess the impact of landmark proximity and shift on accuracy (OE) and precision (ellipse areas), as well as their interaction with instruction (*EGO/ALLO*) and visual field of response (Table 3). To treat the PRO and ANTI-task data on a similar footing, we used separate predictors for these tasks.

**Table 3.** A: Fixed Effects of Multi-level Model using Ellipse Areas in ALLO Instruction conditions as the Criterion

| Predictor | *B* | *B*<br>95 % CI<br>[LL, UL] | *t value* | *p* |
| --- | --- | --- | --- | --- |
| (Intercept) | 5.79** | [5.49, 6.08] | 39.33 | < 2.0 e -16** |
| Landmark Shift | 1.11 | [-0.10, 1.44] | 1.07 | 0.29 |
| Target-Landmark Distance | -3.88** | [-4.40, -3.69] | -6.23 | 7.9 e -10** |
| Landmark Shift x Task (*PRO*) | 2.29* | [0.19, 0.36] | 2.19 | 0.03* |
| Target-Landmark Distance x Task (*PRO*) | 0.68 | [0.30, 1.12] | 1.07 | 0.29 |

| Predictor | *B* | *B*<br>95 % CI<br>[LL, UL] | *t value* | *p* |
| --- | --- | --- | --- | --- |
| (Intercept) | 1.25** | [1.75, 2.25] | 6.41 | < 2.7 e -10** |
| Landmark Shift | -0.05 | [-0.08, 0.03] | -1.22 | 0.20 |
| Target-Landmark Distance | -1.05 ** | [-3.04, -0.69] | -2.88 | 4.1 e -3** |
| Landmark Shift x Task (*PRO*) | 0.08 | [-0.19, 0.36] | 0.60 | 0.54 |
| Target-Landmark Distance x Task (*PRO*) | 0.10 | [-0.32, 1.10] | 0.35 | 0.74 |

*Note.* *B* represents standardized regression weights. *LL* and *UL* indicate the lower and upper limits of the confidence interval, respectively. A: Model Adjusted $R^2 = 0.18^*$ , 95 % CI [0.15,0.21]. B: Model Adjusted $R^2 = 0.09^*$ , 95 % CI [0.03,0.15]
\* indicates $p < 0.05$ . \*\* indicates $p < .01$ .

In summary, target-landmark distance and landmark shift amplitude had no significant influence on either reach precision or accuracy in the *EGO-Instruction* condition. In the *ALLO Condition,* target-landmark distance had a significant influence on both precision (p = 7.9 e-10) and accuracy (p = 4.1 e-3), when separate shift amplitudes are included in the model. Specifically, performance was improved at the largest (3 degrees) distance from the target (or its mirror opposite). Finally, an interaction with Task was also observed (p = 0.03), where the *ANTI Task* showed a stronger association between the degree of landmark shift and precision, compared with the *PRO Task*.

### 3.3 Gaze-Centred Overshoot

One potential source of error is the phenomenon of gaze-centered overshoot, sometimes called ‘retinal magnification’ (Bock, 1986; Enright, 1995; Henriques et al., 1998). In this case, the overshoot is expected to be ‘outward’ relative to the central fixation point. Fig. 7 shows the data distributions of overshoot errors (across participant means) for both the *EGO-Instruction* (blue) and *ALLO-Instruction* data, separated into LEFT and RIGHT visual fields. Overall, the data showed the expected trends, i.e., leftward overshoots in the left visual field and rightward overshoots in the right visual field, but the magnitudes were smaller for the *ALLO-Instruction*, especially in the *ANTI-Task* (Fig. 8 B).

**Fig. 7:**
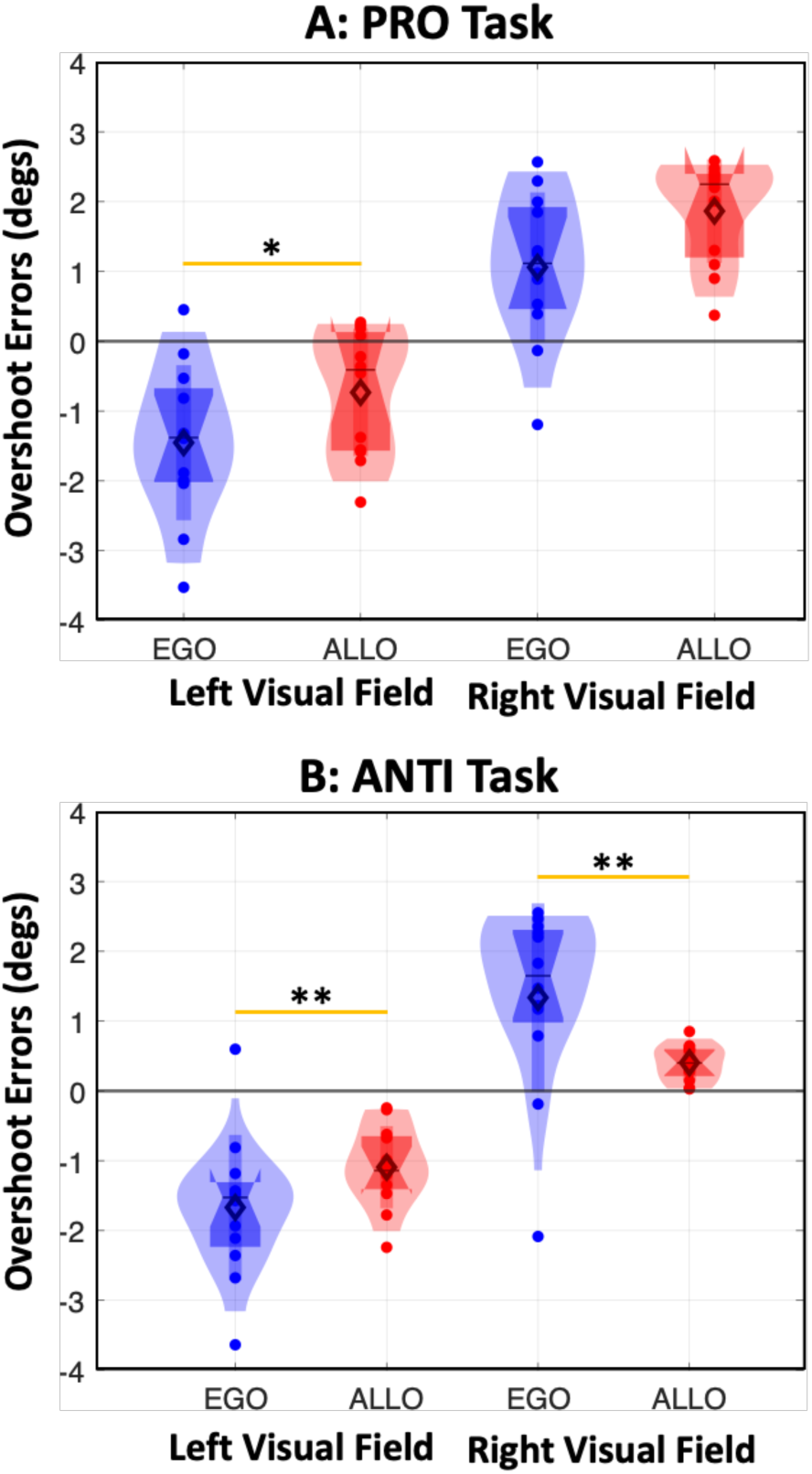
Mean Overshoot Errors for *EGO Instruction* (blue) and *ALLO Instruction* (red) data. (same graphic conventions as Fig. 5). To obtain each data point, the horizontal distance between reach endpoints and expected target location (relative to fixation) was averaged separately for each goal position and then across goals within the left and right visual fields respectively. **(A)** *PRO Task* data. **(B)** *ANTI-Task* data. Significant differences are shown by an asterisk, ‘*’ p < 0.05, ‘**’ p < 0.01.

**Fig. 8:**
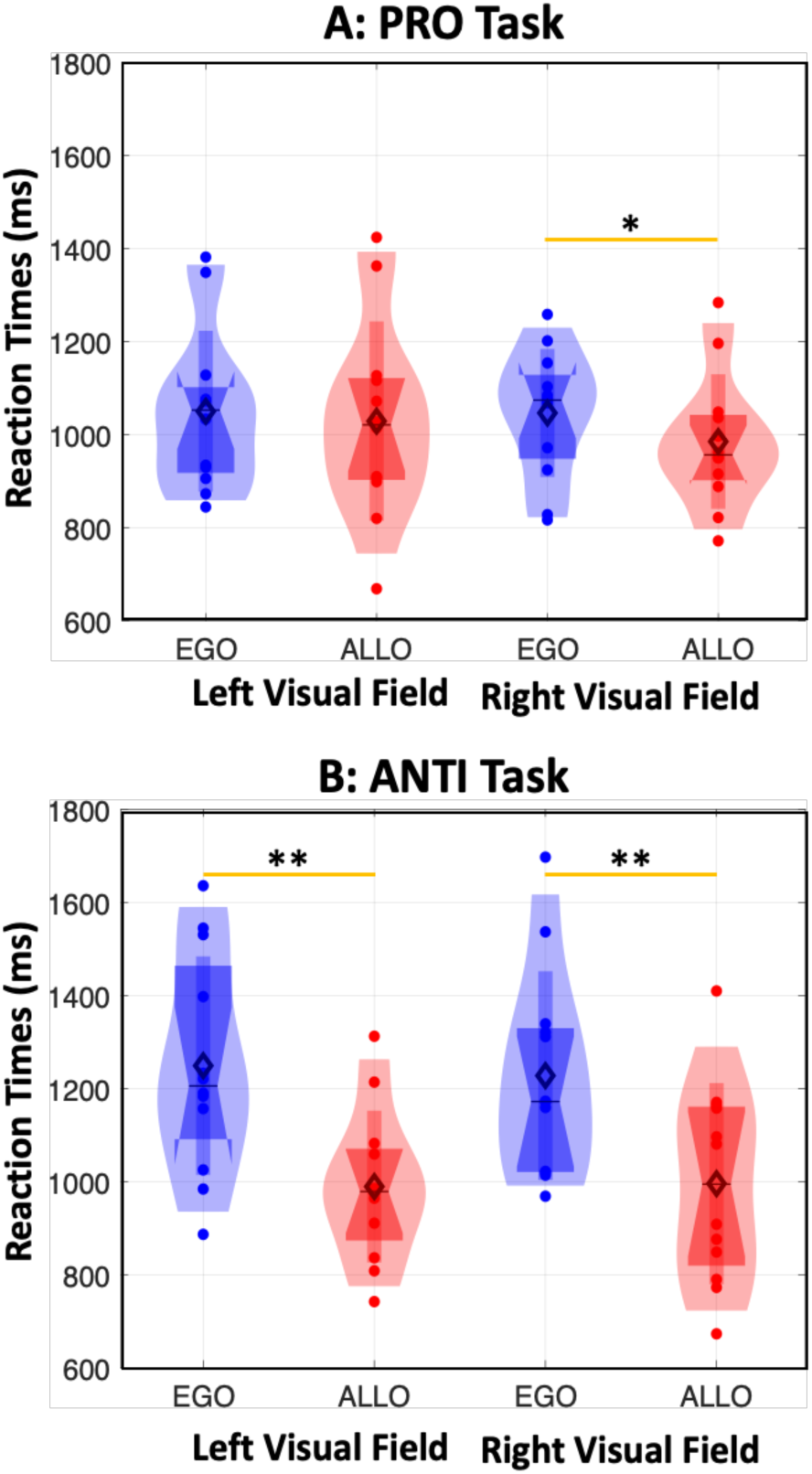
Mean Reaction Times (same graphic conventions as Fig. 5). The reaction times were averaged for targets on the right and left visual fields of response (right and left side of the figure, respectively). **(A)** In the *PRO-Task*. **(B)** In the *ANTI Task* data. *EGO* and ALLO *Instruction* conditions are shown as blue and red in the figure, respectively. Significant differences are shown by an asterisk, ‘*’ p < 0.05, ‘**’ p < 0.01

We analyzed the influence of instruction, task, and visual field on gaze-centered overshoots using a generalized linear mixed model (Table 4). This analysis revealed a significant effect of *EGO/ALLO Instruction* (p = 8.9 e-9) and interaction effects which were analysed using post-hoc t-tests. The effect of *EGO/ALLO* instructions was conditional, depending on the visual field of response and whether participants pointed to the same or opposite side of the target. In the *PRO task*, the *ALLO-Instruction* only produced significantly smaller overshoot errors the left visual field (p = 0.03), whereas it produced significantly smaller errors in both the left (p = 3.1 e-4) and right (p = 2.7 e-4) visual fields in the *ANTI-Task*. The second interaction indicated a significantly higher overshot error in the *PRO-Task* compared with the *ANTI-Task*, but only in the right visual field. Notably, this effect was solely driven by overshoot reaching errors in the *ALLO Instruction* condition, as the absolute overshoot error was less in the *EGO Instruction* condition when participants had to pro-point rather than anti-point. Overall, the main finding here was that the *ALLO Instruction* appeared to suppress gaze-centered overshoot errors in these tasks.

**Table 4.** Fixed Effects of Multi-level Model using Absolute Overshoot Error as the Criterion.

| Predictor | *B* | *B*<br>95 % CI<br>[LL, UL] | *t value* | *p* |
| --- | --- | --- | --- | --- |
| (Intercept) | 2.00** | [1.75, 2.25] | 15.69 | < 2.0 e -16** |
| Task (*PRO*) | -0.17 | [-0.44, 0.10] | -1.23 | 0.21 |
| Instruction (*EGO*) | -1.05 ** | [-1.40, -0.69] | -5.82 | 8.9 e -9** |
| Visual Field (RIGHT) | 0.08 | [-0.19, 0.36] | 0.60 | 0.54 |
| Task<br>(*PRO*) x Instruction<br>(*EGO*) | -0.71** | [-1.10, -0.32] | -3.60 | 3.5e -4** |
| Visual Field (RIGHT) x<br>Task x Instruction (*EGO*) | 0.19* | [0.09, 0.49] | 1.67 | 0.04* |

*Note.* *B* represents standardized regression weights. *LL* and *UL* indicate the lower and upper limits of the confidence interval, respectively. Model Adjusted $R^2 = 0.33^*$ , 95 % CI [0.25,0.41] \* indicates $p < 0.05$ . \*\* indicates $p < .01$ .

### 3.4 Reaction Time

Reaction times were assessed from the moment of the GO signal until the movement onset. In general, the *ALLO Instruction* data showed reduced reaction times relative to the *EGO Instruction* data. This advantage was modest in the *PRO-Task* data (Fig. 8 A) but much stronger in the *ANTI-Task* data (Fig. 8 B), where *EGO Instruction* reaction times appeared to be elevated relative to the *PRO*-data.

This was assessed for the 2 (*EGO/ALLO*) conditions x 2 (*PRO/ANTI*) Tasks and 2 (*LEFT/RIGHT*) visual fields using a generalized linear mixed model. The simplified model in Table 4 was fitted as there was no significant three-way interaction effect. A significant effect of *EGO/ALLO Instruction* was observed (P = 4.9 e-4). Participants exhibited significantly slower reaction times in the *EGO Instruction* condition, with a relative delay of 242.75 ms, 95 % CI [109.30, 376.21]. Additionally, a significant interaction effect was identified (p = 9.8 e-3), which was analyzed using a post-hoc interaction contrast: when averaging over the visual field of response, the impact of *EGO/ALLO Instructions* conditions depended on the Task requirements (i.e., whether participants had to point to the same or opposite side from the target). *ALLO Instruction* resulted in faster reaction times, although the effect was only significant when participants had to point in the opposite visual field (p = 0.03), with a reduction of 245.90 ms, 95 % CI [136.9, 355.0]. A similar analysis showed no significant difference in movement duration.

**Table 5.** Fixed Effects of Multi-level Model using Reaction Time as the Criterion.

| Predictor | *B* | *B*<br>95 % CI<br>[LL, UL] | *t value* | *p* |
| --- | --- | --- | --- | --- |
| (Intercept) | 1001.82** | [907.45, 1096.19] | 21.09 | < 2.0 e -16** |
| Task (*PRO*) | 14.30 | [-94.67, 123.26] | 0.26 | 0.79 |
| Instruction (*EGO*) | -242.75 ** | [-376.21, -109.30] | -3.61 | 4.9 e -4** |
| Visual Field (RIGHT) | 18.77 | [-127.73, 90.20] | 0.34 | 0.73 |
| Task<br>(*PRO*) x Instruction (*EGO*) | -204.65 | [-358.75, -50.55] | -2.64 | 9.8 e -3** |
| Visual Field (RIGHT) x<br>Instruction (*EGO*) | -6.32 | [-147.78, 160.42] | -0.08 | 0.94 |

*Note.* *B* represents standardized regression weights. *LL* and *UL* indicate the lower and upper limits of the confidence interval, respectively. Model Adjusted $R^2 = 0.22^*$ , 95 % CI [0.18,0.26] \* indicates $p < 0.05$ . \*\* indicates $p < .01$ .

## 4. Discussion

Previous studies have reported the sensory influence of a visual landmark on reaching behavior (Krigolson & Heath, 2004; Lemay et al., 2004; Obhi & Goodale, 2005; Schütz et al., 2013), or the weighting of egocentric and allocentric cues (e.g. Byrne & Crawford, 2010; Fiehler et al., 2014) but here we provide a detailed behavioral analysis of the specific influence of *instructions* on performance in an otherwise stimulus-matched, cue-conflict task. Specifically, we measured how the instruction to use (or ignore) landmark-centered coordinates influences the accuracy, precision, and timing of memory-guided reaches, either to the same or opposite visual hemifield as the target. Our results show that participants were generally more accurate, precise, and quicker to react in the allocentric instruction condition, especially when reaching to the visual field opposite to the target. We also observed a left / right visual field effect, where performance was worse overall in the right visual field. We will interpret each of these findings and other details below, in terms of previous literature, potential physiological mechanisms, and their practical implications.

### 4.1 Egocentric vs. Allocentric Aiming: Natural vs. Experimental Conditions

Before discussing our data, we are obliged to consider how our experimental design relates to natural reaching behavior (Fooken et al., 2023). First, the normal visual feedback of the goal and hand was removed throughout the memory delay and reach. Reaches based on initial sensory conditions are thought to rely on internal models of the eye-head-hand system (Blohm & Crawford, 2007; Blohm et al., 2008). In general, this provides accurate behavior (Vercher et al., 1994), but with certain errors variable and systematic errors (Henriques et al., 1998; Medendorp & Crawford, 2002; Van Pelt & Medendorp, 2007). It is noteworthy that even without an instruction, the presence of a visual landmark tends to dampen these errors (Krigolson & Heath, 2004; Lemay et al., 2004; Obhi & Goodale, 2005; Schütz et al., 2013), especially as the memory delay increases (Chen et al., 2011).

Second, egocentric and allocentric cues normally complement each other, whereas we introduced a conflict by surreptitiously shifting the landmark during the memory delay. It has been argued that this simulates situations where visual landmarks are unstable (Byrne & Crawford, 2010) or egocentric signals are unreliable (Byrne et al., 2010; Chen et al., 2011). In the absence of explicit instruction, healthy individuals tend to optimally weigh egocentric vs. allocentric cues based on their relative reliability and perceived stability (Byrne & Crawford, 2010). This weighting is remarkably consistent (∼1/3 allocentric, ∼2/3 egocentric) and generalizes well to naturalistic situations, although the exact ego / allocentric ratio is modulated by various factors such as distance and similarity between the target and landmark (Fiehler et al., 2014; Klinghammer et al., 2017). Although we instructed our participants to shift this weighting completely toward one cue or the other, we cannot assume that the other cue had no implicit influence.

Finally, unlike most natural reaches (which employ the implicit processes above) we explicitly required our participants to follow two different rules (egocentric vs. allocentric) based on a verbal instruction and a set of color-coded cues. Specifically, the allocentric instruction required participants to both attend to the landmark and remember its spatial relationship to the target. This required the integration of bottom-up perception, top-down cognition, and environmental factors (Caduff & Timpf, 2008). This additional degree of cognitive processing could interfere with reaching performance. Thus, one cannot assume either that the allocentric instruction would improve performance, or conversely, that the landmark would have no influence in the Egocentric condition.

### 4.2 The EGO Instruction Condition as a Control: General Observations

Our *EGO Instruction* condition was designed as a control for the ALLO condition, based on several assumptions confirmed in our results. First, as expected from previous studies (Goodale & Milner, 1992; Hu et al., 1999; McIntyre et al., 1997; Westwood & Heath, 2003) participants were able to perform the task, i.e., reach positions correlated with the required goal positions, but with considerable variable and systematic errors. In particular, the EGO Instruction data showed the expected gaze-centered overshoots (Henriques & Crawford, 2000; Henriques et al., 1998; Van Pelt & Medendorp, 2007), which likely originate in the comparison between hand and target position signals used to compute the reach vector in visual coordinates (Dessing et al., 2009).

In the *ANTI-task* version of the *EGO condition*, participants had to suppress the normal pro-reach, and then calculate an opposite reach goal in visual coordinates (Cappadocia et al., 2017; Everling & Munoz, 2000). As predicted, this appeared to produce additional egocentric noise in the system, increasing both systematic and variable errors increased in *the ANTI* version of the *EGO* condition.

Finally, neither target-landmark distance nor landmark shift amplitude had a significant influence on performance in the *EGO Instruction* task. This result is consistent with active suppression of landmark information (although we cannot quantify this without a ‘no instruction’ control). Overall, the *EGO Instruction* produced the expected results, providing a control for comparison with the *ALLO Instruction*. We will focus on the difference between these conditions below.

### 4.3 Influence of the Landmark Instruction: Precision and Accuracy

Our main result was that the instruction to reach relative to the landmark generally enhanced both accuracy and precision relative to egocentric instruction. In particular, there was a systematic reduction of the gaze-centered overshoot effect that was observed in our egocentric data and many previous studies (Henriques et al., 1998; Henriques & Crawford, 2000; Van Pelt & Medendorp, 2007). We also found significant improvements in variable error. Likely this advantage was due to the greater attentional weighting on landmark cues, which tend to improve performance (Lemay et al., 2004), especially after a delay (Chen et al., 2011; Goodale & Milner, 1992; Hay & Redon, 2006; Hu et al., 1999; Krigolson & Heath, 2004; McIntyre et al., 1997; Milner & Goodale, 2006).

Although we equalized the stimuli in conditions, our instructions may have interacted with stimulus perception in some way. In contrast to previous studies (Aagten-Murphy and Bays, 2019; Fiehler et al., 2014), landmark influences on performance increased with distance from the target. This is likely because target-landmark difference used here was small (1.15-3.44 degrees) compared to those previous studies, and relative to the distance of stimuli from the fovea 4.57-15.07. The beneficial influence of our nearest landmarks might have been negated by sensory confusion within response fields during initial perception (Ransom-Hogg & Spillmann, 1980) or more likely, confusion between their representations at the time of response. It is also possible that our choice of color cues interacted with the instruction in some fashion (Nakshian, 1964; Nathans, 1999). But overall, these factors do not negate or explain the benefits of instruction, especially in light of the known advantages of landmark-centered coding for spatial memory (Chen et al., 2011; Lemay et al., 2004).

Finally, we cannot find previous literature that focused on the influence of ego / allocentric *instruction* on behavior, but other studies have reported conflicting behavior data from tasks that employed such instructions. For example, Byrne & Crawford (2010) reported that their control egocentric and allocentric reach tasks did not show significantly different variable error. However, there was no cue conflict in their control tasks, and stimuli were not held constant in these tasks (i.e., landmark present in the landmark task but not in the egocentric task). Our results also differed from those of Thaler & Tod (2009), who found reaching precision to be lowest in their allocentric task. But again, there was no cue-conflict in their paradigm, and the task involved hand alignment, which is thought to employ different mechanisms than reach (Goodale et al., 1994; Monaco et al., 2024).

### 4.4 Results Specific to Anti-Pointing: Accuracy and Timing

Our secondary finding was that the advantage of the allocentric instruction was more pronounced for eaches to the opposite visual field than the target, in terms of accuracy, precision and reaction time. As expected, these effects appear to be driven by decreased performance (including larger gaze-entered overshoots) in the *EGO/ANTI* condition relative to the *EGO/PRO* condition, whereas the *LLO* condition showed similar performance for both the *ANTI* and *PRO* task versions. This is likely ecause anti-reaching requires specific cognitive transformations and neural mechanisms Fernandez-Ruiz et al., 2007; Fischer & Weber, 1992; Gail et al., 2009), whereas the mechanisms for eaching relative to the landmark (Chen et al., 2014). However, anti-movements require the reversal f a motor vector plan. It is noteworthy that anti-movements fall within the class of ‘non-standard’ ule-based spatial transformations that pervade much of our modern life (Sergio et al., 2009), so these ehaviors might benefit the most from the presence and use of allocentric landmarks.

### 4.5 Visual Field Dependence

Our tertiary observation was that the benefit of the allocentric instruction for anti-reaching was symmetric across the visual hemifields. Whereas the egocentric anti-pointing task produced ymmetric gaze-centered errors, the allocentric instruction only mitigated these errors for goals / andmarks that shifted from the right to the left visual field. Byrne et al. (2010) found that allocentric andmarks had more influence on pointing when saccades caused remembered targets to remap from he right to the left hemifield. This might be related somehow to a tendency of our participants to be ight-hand and right-eye dominant. Also, various studies have suggested that the right cortical emisphere (corresponding to the left visual) plays a stronger role in allocentric processing than the eft (Faillenot et al., 1999; Galati et al., 2000; Zaehle et al., 2007; Fink et al., 2003; Weiss et al., 2006).

### 4.6 Possible Physiological Mechanisms

Most sensorimotor neuroscience studies only consider the egocentric transformation from sensory to motor coordinates, but to interpret the current results we must consider both 1) the integration of sensory input to construct two different spatial rules, and 2) the implementation of these rules through separate allocentric and egocentric transformations.

For the first ‘instruction’ component, one might safely assume that the auditory instruction is processed through the auditory cortex and higher-level language areas (Friederici et al, 2000; Humphries et al., 2001; Vandenberghe et al., 2002; Wernicke, 1874), whereas initial color and spatial information from the visual stimuli are processed by well-known pathways in the occipital cortex (DeYoe & Van Essen, 1988; Livingstone & Hubel, 1988; Zeki, 1993). The key question is, how does the former impose a rule on how the latter is processed? One likely candidate area is the prefrontal cortex. For example, the lateral prefrontal cortex receives complex multisensory inputs and is thought to play roles in imposing rules on sensory inputs (Miller & Cohen, 2001), including both inhibiting incorrect responses (DeSouza et al., 2003) and selecting correct responses (Rowe et al., 2000). It is also likely that the parietofrontal attention network is involved in directing attention to the landmark (Buschman & Miller, 2007; Bisley & Goldberg, 2003). Consistent with these speculations, the parietofrontal cortex is thought to exert recurrent influence on the early visual areas during reach (Cappadocia et al., 2017; Blohm et al., 2019) and appears to alter visual processing in these areas (Monaco et al., 2024; Velji-Ibrahim et al., 2018).

Once such rules are enacted, it is thought that egocentric and allocentric transformations for reach are processed through separate pathways. Classic neuropsychology experiments suggest that the dorsal visual stream (via parietal cortex) handles egocentric visual transformations (Carey et al., 2006; Goodale & Milner 1992; Schenk, 2006). Numerous studies have examined the role of parietal cortex in action; too many to review here (e.g., Buneo & Andersen, 2006; Crawford et al., 2011; Gallivan & Culham, 2015; Vesia & Crawford, 2012). However, it is pertinent that damage to the dorsal stream alters the gaze-centred errors reported here (Khan et al., 2005a, 2005b), and that the visual field effects observed here could be attributed to interactions between hand and hemifield lateralization in parietal cortex (Medendorp et al., 2005; Perenin & Vighetto, 1988; Rossetti et al., 2003). Parietal cortex also plays a role in coding rule-based egocentric transformations, such as the anti-reach task (Gail & Andersen, 2006; Cappadocia et al., 2017).

In contrast, the ventral visual stream (via temporal cortex) is thought to handle allocentric transformations (Goodale & Milner 1992; Carey et al., 2006; Schenk, 2006). This distinction is supported by neuroimaging experiments that showed spatial tuning for egocentric targets in the dorsal stream and allocentrically defined reach targets in the ventral stream (Chen et al., 2014). Ultimately, the latter must be integrated into the motor system for egocentric control of action, and this appears to happen in both the parietal and frontal cortex (Chen et al., 2018). Consistent with this, frontal cortex visual responses integrate target and landmark location (Schütz et al., 2023), whereas their memory and motor responses for gaze are able to integrate allocentric and egocentric coordinates (Bharmauria et al., 2020, 2021) in a cue-conflict task similar to that used in reach studies.

## 5. Conclusions

Whereas several studies have reported the influence of allocentric landmarks on reach, and some studies employed explicit instructions to reach in allocentric coordinates, this study examined the interaction of these two factors: whether an explicit instruction to attend to and use a landmark to encode reach direction provides additional behavioral benefits. We found that reach performance (accuracy, precision, reaction time) was generally enhanced, especially in the more difficult task of reaching toward a location defined in the visual hemifield opposite to the target. This has the practical implication that explicit instructions to use visual landmark cues may enhance performance, especially in high-demand ‘non-standard’ spatial tasks (Sergio et al., 2009; Dalecki et al., 2019), and for individuals with degraded egocentric transformations due to age, developmental disorders, or brain damage (e.g., Goodale et al., 1994; Khan et al., 2005; Niechwiej-Szwedo et al., 2011; Tippett et al., 2007).

## 6. Acknowledgements

The authors thank V. Bharmauria for helpful comments and proof reading, and S. Sun for assistance with coding.

